# Molecular Engineering of Silk via Fatty Acid Esterification for Water-Stable, Anti-fouling, Biocompatible, Implantable Materials

**DOI:** 10.64898/2026.10.20.760528

**Authors:** Sanjana Gopalakrishnan, Emilee Zhang, Mariah L. Arral, Edward B. Gordon, Sophie M. Steedly, Jugal K. Sahoo, Chunmei Li, Weiguo Hu, David L. Kaplan

**Affiliations:** Department of Biomedical Engineering, School of Engineering, Tufts University, Medford, MA 02155; Department of Polymer Science and Engineering, College of Natural Sciences, University of Massachusetts Amherst, Amherst, MA 01003

## Abstract

Biopolymers offer attractive properties for implantable materials, but their limited aqueous and enzymatic stability constrains long-term performance in biological environments. Silk fibroin is mechanically tunable and thermoplastically processable, yet remains susceptible to swelling in aqueous environments, which can impact mechanical properties. Here, we establish fatty acid esterification as a strategy to transform silk into a hydrophobic, water-stable, yet still bioresorbable material (FAsilk). Fatty acid functionalization significantly increased resistance to aqueous swelling and enzymatic degradation while preserving structural and mechanical integrity and thermoplastic processability. The resulting FAsilk biomaterials also reduced microbial contamination, blood clot formation, and macrophage adhesion relative to native silk, due to the change in hydrophobicity, while maintaining cytocompatibility and hemocompatibility. Subcutaneous implantation in mice demonstrated favorable biological responses, including minimal impacton body weight and no significant pro-inflammatory responses, indicating no adverse effects on animal health. Thus, fatty acid functionalization enables silk to combine hydrophobicity and aqueous stability with processability and biodegradability, providing a mechanically robust materials platform for bioresorbable implants, including medical devices, tissue-engineered grafts and drug delivery systems.

## Introduction

The significant advantages of biopolymers over conventional synthetic polymers^1^ have increasingly led to their adaptation in biomedical applications such as tissue engineering,^2^ drug delivery,^3^ and regenerative medicine.^4^ Biopolymers are naturally abundant, biocompatible, and bioresorbable which make them good candidates for implantable materials.^5^ However, their performance in physiological environments is limited by their poor aqueous stability, high propensity for microbial contamination, and poor hemocompatibility^.6,^ Biopolymers are prone to uncontrolled hydrolytic and proteolytic degradation, resulting in structurally and mechanically weak implants.^7^ Furthermore, interactions between some biopolymers and blood can promote hemolysis, protein adsorption, and platelet activation, while their susceptibility to biological fouling and microbial adhesion can increase the risk of thrombosis and implant-associated infection.^8^ These limitations remain major barriers to the broader application of biopolymers as implantable materials, including in devices such as catheters, vascular grafts, and surgical implants, due to associated risks of device failure.

Silk fibroin (hereafter referred to as silk) is a naturally abundant protein extracted from the *Bombyx mori* silk cocoons and has distinct advantages over other biopolymers for biomedical applications.^9^ Silk exhibits exceptional mechanical tunability, relatively slow biodegradation in vivo due to proteolysis, in addition to hemocompatibility, making it useful for implantable biomater ials.^10^ Furthermore, silk protein can be processed at high temperatures and pressures, leading to machinable, dense and hard implantable materials with high mechanical strength.^11^ However, poor aqueous stability due to increased water uptake remains a critical challenge with silk-based implants. Furthermore, native silk, similar to other biopolymers, is prone to microbial contamination which is a leading cause of implant failure.^12^ In a recent study, we developed hydrophobic silk materials by tethering perfluorocarbons to the silk backbone^13^ and demonstrated its ability to minimize microbial contamination and platelet activation,^14^ however, the use of perfluorocarbons and consequent exposure to (polyfluoroalkyl substances (PFAS) poses potential environmental and health concerns.

In the present study, we developed a novel approach for fatty acid modification of silk (FASilk) via Steglich esterification. We conduct esterification^15^ in a silk ionic liquid to access the active amino acid residues buried within the β-sheet structures, thereby achieving a high degree of functionalization while preserving favorable properties of native silk. We demonstrate that this approach is versatile and can be used to modify silk with different fatty acids to impart functional properties to FAsilk. Importantly, we successfully tethered α-linolenic acid, a functional component of omega-3 fatty acids known to support vascular health,^16^ demonstrating the utility of this approach for biomedical applications. We establish that FAsilk thermoplastics successfully minimize water uptake and, thereby crucially, improve mechanical and structural stability in aqueous environments compared to native silk, while remaining biodegradable. Lastly, we establish the biocompatibility of FAsilk post-chemical modifications through *in vitro* and *in vivo* evaluations. FAsilk remains hemocompatible, non-immunogenic, and non-toxic, as observed through our *in vitro* and *in vivo* studies. FAsilk exhibits comparable biocompatibility to native silk when implanted in animals over seven days based on both histological analysis of implant site tissue and inflammation panels. Importantly, FAsilk demonstrated significant improvements in anti-fouling behavior compared to polytetrafluoroethylene (PTFE) and native silk against *S. aureus* biofilms. Together, these findings establish FASilk as a promising platform for implantable biomaterials and medical devices combining aqueous stability, biocompatibility, and antifouling properties.

## Results and Discussion

### Fatty Acid Modification of Silk

As illustrated in Figure 1(a) the protein structure of silk contains significant portions of β-sheets (crystalline domains) connected together by less organized linkers.^17^ The predominant amino acid residues within the silk β-sheets are glycine, alanine and serine,^18^ with the serine residues amenable to chemical modification butsterically inaccessible.^19^ We therefore utilized BMIM-Cl (1-butyl-3-methylimidazolium chloride) to prepare silk ionic liquids to disrupt these crystalline domains to increase accessibility of serine residues for modification.^20^ Additionally, threonine and tyrosine residues present in silk fibroin in lower quantities are also amenable to esterification, though degree of modification is expected to be significantly lower than serines. Fatty acid modifications were conducted using different fatty acids for reactions on the serine residues via Steglich esterification (Figure 1b, Figure S1, Figure S2). Furthermore, the degree of disruption was enhanced by including 8M urea to enable a higher degree of substitution.^21^ Low- and high-modified silks were prepared with and without the inclusion of urea respectively (Figure S3). As seen in Figure S1, several peaks corresponding to the fatty acid were observed in both the modified silks. Additionally, Figure S3 showed that peaks corresponding to the carbonyl bonds and alkoxy bonds are present in modified silks but absent in native silk, indicating successful ester bond formation (C=O at 1700 and 1740 cm^-1^; C-O at 1170 and 1230 cm^-1^). Importantly, an increased absorbance was observed in the carbonyl peaks in high-FA compared to low-FA, consistent with a higher degree of substitution as expected from the inclusion of urea.^21^

**Figure 1.**
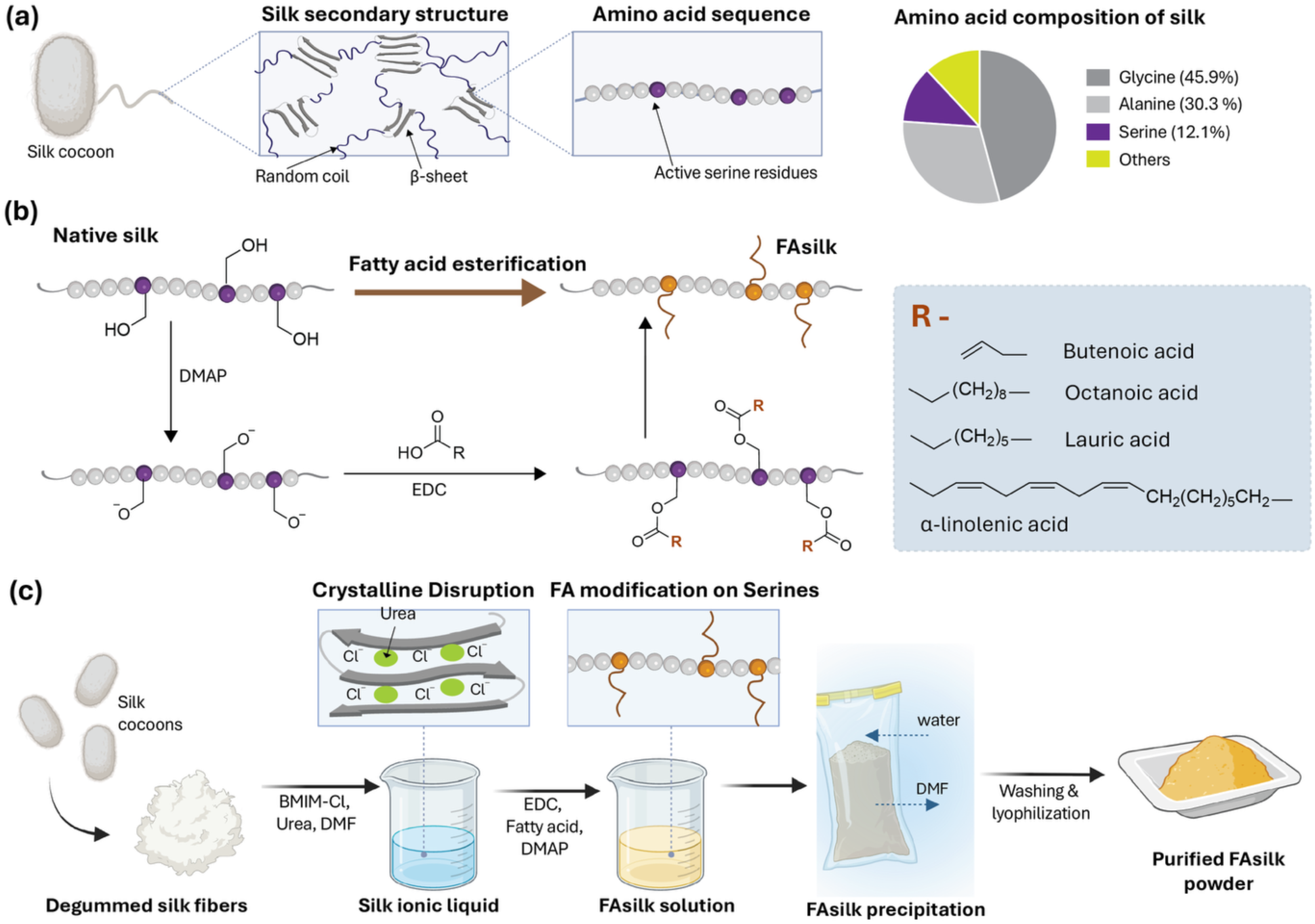
Chemical modification and processing approach to prepare FAsilk materials. (a) Protein structure of silk containing β-sheets and random coils, as well as the amino acid sequence and composition. Silk contains 12.1% serine residues, which are utilized for esterification in addition to threonine and tyrosine. (b) Steglich esterification of serine dues, which are deprotonated by DMAP, while EDC results in activated fatty acid esters. When combined, fatty acids are tethered to the serine residues via esterification leading to FAsilk. FAsilk was prepared using four different fatty acids, illustrating the versatility of this approach. (c) Process of synthesis of FAsilk involves first preparing silk ionic liquids with or without urea to improve accessibility to serine residues in the β-sheet, followed by esterification of the serine residues, followed by purification via precipitation in water. FAsilk precipitate is further washed and dried to obtain purified FAsilk powder.

Briefly, degummed silk fibers were melted in BMIM-Cl (1-butyl-3-methylimidazolium chloride) to produce a silk ionic liquid (SF_IL_). SF_IL_ was then diluted in dimethylformamide (DMF) containing DMAP (4-Dimethylaminopyridine), EDC (N-Ethyl-N′-(3-dimethylaminopropyl)carbodiimide) and the fatty acid of choice. To achieve a high degree of modification, 8 M urea was included in the above reaction mixture. The reaction was stirred at room temperature for 24 h and dialyzed against deionized water in a 3.5 kDa dialysis tubing for 72 h, with 6 periodic water changes, and at the end of dialysis, the modified silk precipitated from the reaction mixture due to low water solubility. The precipitated FAsilk was collected and further washed through continuous stirring in 500 mL of deionized water for an additional 48 h with multiple water changes. The purified FAsilk was frozen at -80°C, lyophilized and stored at -20°C. Figure 1(c) depicts the process of synthesis and purification of FAsilk. To assess the versatility of this approach, we modified silk fibroin with fatty acids of different chain lengths (Figure S2). Additionally, we tested the water uptake of both lauric acid- and α-linolenic acid modified silk plastics (Figure S5) and noted that α-linolenic acid-modified plastics performed significantly better than lauric acid, likely due to the longer chain length (C18 vs C12) and presence of an unsaturated region (3 alkene bond for α-linolenic acid in comparison to none for lauric acid). Therefore, we selected α-linolenic acid for the remainder of our studies due to its improved aqueous stability, established anti-inflammatory properties, and potential biomedical applications^16^ (hereafter referred to as FAsilk).

### Characterizing the Structural and Mechanical Properties of Fatty Acid Modified Silks

Implantable materials have previously been fabricated from silk by thermopressing into dense, hard plastics, that can be molded or machined into different shapes.^11^ Here we studied the process of thermal pressing of the FAsilks and the role of the chemical modification on the molecular rearrangement, and consequently structural and mechanical properties, with a particular focus on swelling in water related to changes in mechanical properties.

The process of thermopressing native silk typically results in the reorganization of the protein structure facilitated by hydrogen bonding interactions in the flexible regions and unbound water molecules,^22^ leading to increased crystallinity and a higher content of β-sheet (Figure 2a,b). As seen in the secondary structure composition obtained from ATR-FTIR measurements in Figure 2(b) and Figure S3, native silk powders contain a low percentage of β-sheet, and a higher percentage of random coil and α-helices. During thermopressing, there is a significant increase in percent β-sheet when compared to the native silk powder, and a significant reduction in the percentage of random coil and α-helix (Figure 2b). In contrast, with this new modified silk, FAsilk, the protein secondary structure remained constant with a high percent β-sheet content relative to other secondary structures. Ordinary one-way ANOVA showed a statistically significant effect of the type of silk on the percent β-sheet content in the materials (powders, dense plastics) [F(5,12) = 54.59, p < 0.0001], random coil [F(5,12) = 22.23, p < 0.0001], α-helix [F(5,12) = 31.55, p < 0.0001] and β-turns [F(5,12) = 20.52, p < 0.0001]. Post-hoc pairwise Tukey tests showed statistical significance across multiple pairs (Figure 2b). Importantly, while both native silk and FAsilk can be formed into rigid plastics via thermal pressing, lyophilized SF_IL_ did not form into the dense plastics at comparable temperatures, leading to a brittle powder (Figure 2(c)), despite the higher relative percent of β-sheet content when compared to the native silk powder (Figure S3). Therefore, changes to the protein crystallinity and percent β-sheet content alone do not explain the plasticizability of silk plastics.

**Figure 2.**
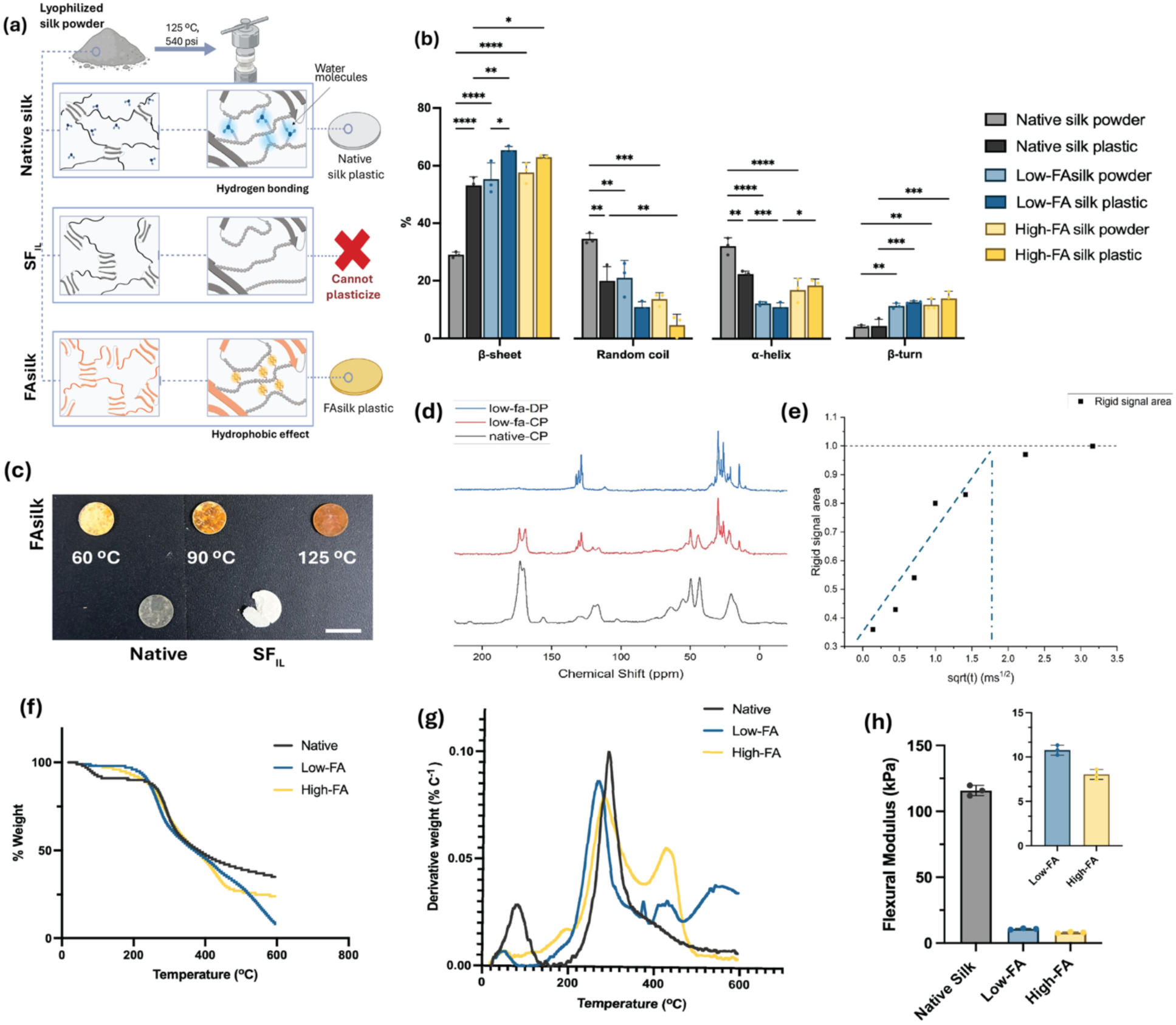
Protein structure remodeling during FAsilk synthesis and impact on plasticization. (a) Schematic illustrating the changes in molecular interactions in native silk, SF_IL_ and FAsilk in lyophilized powder form and when plasticized via thermopressing. (b) Protein secondary structure of native silk, SF_IL_ and FAsilk measured and calculated using ATR-FTIR spectroscopy. (c) Images of native silk plastic, thermopressed SF_IL_, and FAsilk plastics prepared at 60°C, 90°C and 125°C. FAsilk transitions from unplasticized substance (heterogenous, rough) at 60°C to smooth, homogenous plastic at 125°C, while SF_IL_ remains unplasticized at 125°C. (d) 13C MAS NMR spectra showing (blue) DP spectrum of low-FA, with mostly signals from mobile segments; (red) CP spectrum of low-FA, with mostly signals from rigid segments; (black). (d) CP spectrum of native silk sample. (e) Normalized signal area of the rigid segments as a function of the square root of diffusion time. Visually drawn trendline at the early phase of the diffusion crosses the saturation at a sqrt(t) of 1.8 ms1/2. Thermogravimetric analysis showing (e) %weight loss as a function of temperature (°C) and (f) derivative weight peaks as a function of temperature (°C). All silks show similar degradation peaks at ∼200 to 250°C. Additionally, native silk shows a peak at ∼ 100°C, which corresponds to water vapor evaporation. (g) Flexural modulus (kPa) of native silk and FAsilks measured by three-point bending tests, showing that native silk is stronger than FAsilks, presumably due to differences in molecular organization. Asterisks indicate adjusted p-values. p < 0.05 (*), p < 0.01 (**), p < 0.001 (***), and p < 0.0001 (****). Unmarked comparisons are not significant.

Therefore, we concluded that enhanced molecular mobility, in combination with increasing crystalline domains is crucial for thermal fusion during thermopressing of silk. We hypothesized that the fatty acid esterification not only causes increased molecular rearrangement, leading to higher percent β-sheet content, but also facilitates thermal fusion during thermopressing through increased molecular mobility^23^ potentially due to the increased hydrophobic interactions between fatty acid groups. In contrast, the molecular mobility in native silk is facilitated by the presence of unbound water which leads to increased hydrogen bonding interactions between silk chains and enables thermal fusion. Previous studies have also demonstrated that increased water content in silk improved the plasticizability and enabled thermal fusion at lower temperatures, further supporting our hypothesis.^22^

Our hypothesis is further supported by ^1^H and ^13^C solid-state NMR conducted on native and low-FAsilk (Figure 2(c), Figure S4). ¹³C CP/MAS (cross-polarization/ magic-angle spinning) and short-recycle-delay DP/MAS (direct polarization/ magic-angle spinning) NMR spectra support the presence of a mobile FA-rich phase alongside predominantly rigid silk segments. The short-recycle-delay ¹³C DP spectrum (blue) preferentially highlights FA-associated resonances, whereas the CP spectrum (red) retains prominent silk signals, supporting the coexistence of mobile FA-rich regions and predominantly rigid silk segments (Figure 2(d)). Consistently, low-FA silk exhibits a substantially larger narrow ¹H component (∼7 kHz) than native silk, indicating an increased mobile proton population (Figure S4). Furthermore, Silk-associated NMR signals increase as magnetization transfers from mobile FA-rich regions to rigid silk segments (Figure S4). The rapid increase and eventual plateau in signal intensity (Figure 2(e)) indicate that these regions are closely spaced, with an estimated domain size of approximately 3 nm based on the spin-diffusion model. This shows that the linolenic acid domains are finely dispersed throughout FAsilk and may serve a similar role as water in native silk sample, increasing the mobility of the amorphous phase and facilitating the melt-state processing of the FA modified silk.

SF_IL_ prepared under anhydrous conditions lacks the native hydrogen-bonding network and hydrophobic interactions introduced by the fatty-acid functionalization, resulting in insufficient intermolecular cohesion for plasticization. This is further supported by thermogravimetric analysis (Figure 2(f,g)), where a mass degradation peak corresponding to water vapor evaporation was observed at ∼100°C ^24^ in native silk, but was absent in high-FAsilk and substantially smaller in the low-FAsilk, demonstrating that fewer water molecules are present within the FAsilk matrix when compared to native silk.

Next, we evaluated the impact of the secondary structure on mechanical properties of native and FAsilks. As seen in Figure 2(h), there was a significant reduction in flexural modulus of the FAsilks as compared to native silk in the dense plastic format, despite similar percent β-sheet (Figure 2(b)). Ordinary one-way ANOVA showed a statistically significant effect of type of silk on the flexural modulus [F(2, 6) = 2183, p < 0.0001]. Post-hoc pairwise Tukey tests showed statistical significance across all pairwise comparisons. In native silk, thermopressing is accompanied by a substantial reduction in flexible domains in the secondary-structure, such as random coils, resulting in a rigid and mechanically robust plastic. In contrast, FAsilk plastics retained a higher β-turn content relative to native silk plastics, suggesting greater conformational flexibility within the protein network. This difference, together with the altered protein structure resulting from FA functionalization, may contribute to the distinct mechanical properties of FAsilk plastics, including the reduction in flexural modulus.

We therefore concluded that fatty acid modification with silk ionic liquids significantly alters the molecular interactions in modified (FAsilk) silk. However, FAsilk retains thermal plasticization at high temperatures and pressure crucial for the fabrication of bioactive medical devices and implants via the formation of dense, plastic materials.

### Evaluating the Effect of Fatty Acid Modification on Aqueous and Enzymatic Stability

Implantable biomaterials must exhibit exceptional aqueous and enzymatic stability over prolonged incubation periods to minimize the risk of structural distortions, such as swelling, mechanical weakening, and uncontrolled enzymatic degradation.^25^ We tested the aqueous stability of these new FAsilk plastic materials over prolonged incubation in aqueous media by measuring the percent water uptake and volumetric swelling over time. FAsilk plastics were prepared with both low and high degrees of modification with fatty acid, and compared with native silk plastics as control. All samples were incubated in phosphate buffered saline (1X, pH-7.4) for a period of 7 days at 37°C. Percent water uptake was measured against the initial dry weight of each sample and is normalized accordingly, as seen in Figure 3(a,b), both low- and high-modified FAsilk plastics had significantly reduced percent water uptake, as compared to native silk plastics. Ordinary one-way ANOVA showed a statistically significant effect based on the type of silk on the percent of total water uptake [F(2, 9) = 44.58, p < 0.0001]. Post-hoc pairwise Tukey tests showed statistical significance between native silk and both FAsilks (Figure 3b). This was further demonstrated by the change in surface morphology and surface roughness post-7-day incubation (Figure 3c,d). Ordinary two-way ANOVA showed statistically significant hydration [F(1, 24) = 60.64, p < 0.0001] and a significant interaction between hydration and silk type [F(2, 24) = 15.30, p < 0.0001], with no statistically significant effect of the type of silk alone [F(2, 24) = 2.203, p = 0.1323]. Post-hoc pairwise Tukey tests showed statistically significant differences between dry and wet conditions for native silk and low-FAsilk likely due to swelling which impacts surface features, but not for high-FAsilk indicating low swelling in aqueous conditions (Figure 3c, d). We concluded that the hydrophobic modification, particularly the high degree (Figure S5) of FA modification, significantly improved the aqueous stability of silk plastics and the degree of modification played a role in the overall aqueous stability.

**Figure 3.**
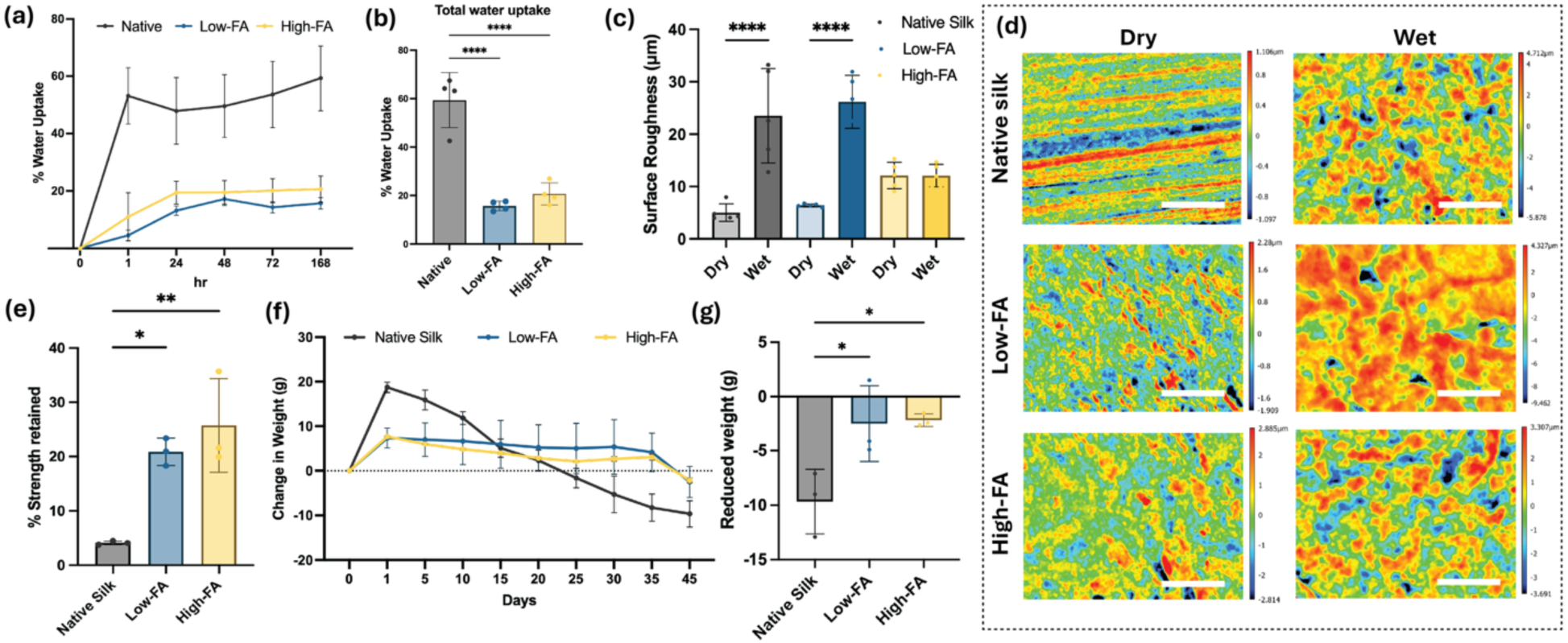
Stability of FAsilk materials. (a) %Water uptake measured by change from initial mass over a period of incubation for 7 days. (b) Total %water uptake after 7 days, normalized against the dry weights of native and FAsilk plastics. Low- and High-FAsilk plastics both showed significantly lower water uptake as compared to native silk. (c) Surface roughness of dry (day 0 of incubation) and wet silk plastics (day 7 of incubation). While native and low-FAsilk showed a significant change in surface roughness, no significant change was observed in high-FA silk plastics. (d) Surface morphology of dry and wet silk plastics, measured by 3D surface profiler. Scale bar is 200 μm. (e) %Strength retained measured by comparing initial flexural modulus of dry samples to final modulus after 7 day incubation. Native silk plastics retained only about 5% of their initial mechanical strength, compared to FAsilk plastics which retained ∼ 20%. (f) %Mass loss measured over time during prolonged incubation in 10 U/mL of Protease XIV. (g) Total %Mass loss after 45 days of enzymatic degradation showed significantly lower mass reduction in FAsilks compared to native silk, indicating improved enzymatic stability. Asterisks indicate adjusted p-values. p < 0.05 (*), p < 0.01 (**), and p < 0.0001 (****). Unmarked comparisons are not significant.

We next characterized the impact of prolonged incubation in an aqueous environment on the mechanical properties of FAsilk plastics by measuring the flexural modulus of wet samples post 7 day aqueous incubation via a three-point bending. While the dry flexural modulus of FAsilk was significantly lower than the native silk plastics (Figure 2f), Figure S6), both low- and high-modified FAsilk retained ∼20% of their dry mechanical properties post-incubation compared with native silk which retained <5% (Figure 3e). Ordinary one-way ANOVA showed a statistically significant effect of type of silk on the percent total water uptake [F(2, 6) = 14.23 p = 0.0053]. Post-hoc pairwise Tukey tests showed statistically significant difference in the percent strength retained between native silk and FAsilks but not between the two FAsilks (Figure 3e). However, high-FAsilk exhibited an overall upward trend in percent strength retained, suggesting that higher hydrophobicity contributed to improved mechanical stability. Therefore, we concluded that simply incorporating fatty acids in the silk backbone minimized the loss of mechanical strength in aqueous environments. Furthermore, it may be possible to improve the absolute mechanical strength of FAsilk materials by altering the molecular weight of the silk prior to the chemical modification and by utilizing post-fabrication crosslinking approaches.

Finally, we characterized the rate of enzymatic degradation of FAsilk plastics as compared to native silk plastics. Low- and high-modified FAsilk plastics, as well as native silk controls, were incubated in 20 U/mL concentration of Protease XIV at 37°C over a period of 30 days, and the change in mass was recorded. Figure S7 also shows the biodegradablilty of FAsilks powders when mixed with protease XIV solution. a A dark smear averaging at ∼110 kDa was observed for Native silk and lighter smears were observed for low-FA and high-FA mixed with water, whereas clear bands at ∼3.5 kDa was observed for all three silks (native, low-FA and high-FA) post incubation in protease solution. This reduction in molecular weight is attributed to the degradation of the protein. As seen in Figure 3(f,g) and Figure S7, while both FAsilk and native silk plastics remained biodegradable, the enzymatic degradation of both FAsilk plastics was significantly slower than native silk, as indicated by the lower reduction in mass over the same period of incubation. Ordinary one-way ANOVA showed statistically significant effect of type of silk on the percent total water uptake [F(2, 6) = 7.605, p = 0.0226]. Post-hoc pairwise Tukey tests showed statistical significance between native silk and both FAsilk plastics, as indicated in Figure 3(g). Taken together, this demonstrated that fatty acid-modification improved enzymatic stability while preserving biodegradability.

We therefore concluded that, fatty acid modification improved aqueous and enzymatic stability of silk by maintaining structural and mechanical properties during prolonged incubation, while still preserving biodegradability.

### Assessing biomaterial interaction with the biological environment

The interaction of biomaterial with biological milieus regulates critical processes including microbial contamination, biofouling, thrombosis and immune responses. ^26^ We hypothesized that fatty acid modification would impart anti-fouling properties to silk due to the increased hydrophobicity, leading to low microbial and cellular adhesion. Since the high-FAsilk samples showed the most significant improvement in aqueous stability (Figure 3), further biological studies focused on studies with the high-FAsilk materials in comparison to the silk control materials.

Uncontrolled adhesion of blood components such as proteins, platelets or cells on the surface of biomaterials leads to clot formation and subsequently thrombosis. We therefore conducted a thrombogenicity assay on the silk biomaterials by evaluating clot formation through standardized protocols.^27^ Briefly, 10 μL of whole human blood was deposited onto each sample to enable clot formation. Samples were then submerged in deionized water to enable lysis of nonadhered RBCs. The extent of clot formation was estimated based on the amount of hemoglobin released into solution due to the lysis of RBCs, with lower absorbance indicating higher retention of blood components on the sample and, therefore, higher clot formation. The absorbance measured for each sample was compared with that of a 100% hemolysis control, prepared by directly incubating 10 μL of whole blood in 500 μL of deionized water and the percentage of clot formation for each sample was calculated as the percentage decrease in absorbance relative to the 100% hemolysis control. As seen in Figure 4(a), both the FA-modified and unmodified silk samples showed comparably low clot formation to the tissue culture plate (TCP) and PTFE controls, indicating minimal adhesion of blood components and consequently a low propensity for thrombogenicity. Ordinary one-way ANOVA showed no statistically significant differences [F(4,10) = 1.470, p = 0.2824]. This finding is in concert with prior studies with silk biomaterials where low thrombogenicity was reported.^28^

**Figure 4.**
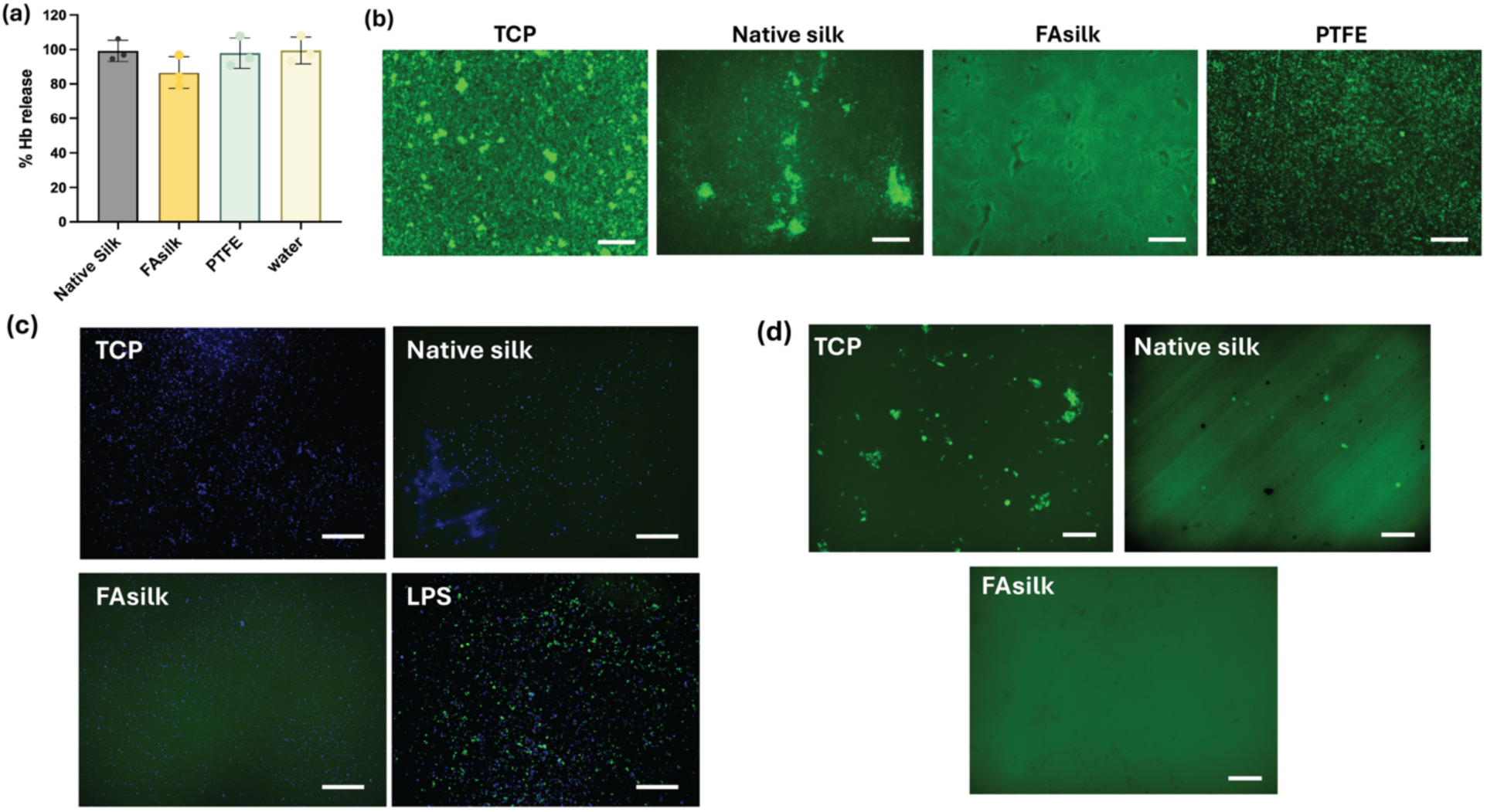
Interaction of biological milieus with FAsilk. (a) Thrombogenicity assessed by blood clotting assay. %Hb (hemoglobin) lysis was inversely proportional to adhesion of blood components and clot formation. (b) Biofilm adhesion of *S.aureus* (green) bacteria showing visibly higher adhesion on TCP and native silk as compared to FAsilk. Scale bar is 200 μm. (c) Monocyte activation tested using THP1-ASC-GFP cells on TCP, Native and FAsilk showed no significant activation (green) as compared to positive control LPS. Nuclei stained using DAPI (blue). Scale bar is 200 μm. (d) Adhesion of differentiated macrophage-like THP1-ASC-GFP cells on TCP, native silk and FAsilk. FAsilk samples showed little to no adhesion of cells. Scale bar is 400 μm.

We next evaluated the propensity for microbial contamination of the silk substrates by challenging them with a high density of *S. aureus* microbes under biofilm forming conditions. Silk plastics were incubated in a high concentration of bacterial solution to encourage biofilm formation. After 24 h, adhered bacteria were stained and imaged via fluorescence microscopy. As seen in Figure 4(b), lower numbers of bacteria are observed on the surface of the FAsilk plastics as compared to native silk and PTFE controls. The diffuse fluorescence observed in both silk samples was due to the auto-fluorescence of silk. Therefore, FAsilk outperformed commercially utilized synthetic materials in preventing microbial fouling.

Finally, we investigated immune cell adhesion to the material surface as an indicator of the potential for a foreign body response. The foreign body response is a localized immune reaction triggered by recognition of a foreign biomaterial, in which circulating monocytes adhere to the material surface and differentiate into macrophages. ^29^ These macrophages can recruit additional immune cells and initiate inflammatory signaling cascades, resulting in sustained local inflammation that can ultimately compromise implant integration and function. Monocyte activation and macrophage recruitment were both independently investigated using THP-1-ASC-GFP reporter cells. To study monocyte activation, unprimed THP-1-ASC-GFP cells were incubated on silk plastics and the GFP expression, which is correlated to NF-κB dependent inflammatory activation. As seen in Figure 4(c), no GFP expression was observed from cells incubated on the silk plastics for 24 hours as compared to the positive control where cells were activated using lipopolysaccharides (LPS). Next, the adhesion of differentiated monocytes was studied. THP-1-ASC-GFP monocytes were first differentiated using PMA (phorbol 12-myristate 13-acetate), which induces macrophage-like polarization and results in diffuse GFP expression.^30^ The differentiated monocytes were then incubated with silk plastics to evaluate macrophage adhesion. As seen in Figure 4(d), few adhered cells were observed on the FAsilk, indicating a significantly reduced likelihood of a foreign body response owing to the fatty acid modifications.

Based on these studies, we concluded that fatty acid-modified silk successfully minimized biofouling, microbial contamination, monocyte activation and macrophage adhesion, supporting its suitability for implantable biomaterials. Importantly, FAsilk significantly reduced microbial contamination and macrophage recruitment as compared to native silk, demonstrating improved biological performance due to fatty acid modification. Furthermore, FAsilk significantly outperformed PTFE in preventing microbial contamination while demonstrating comparably low thrombogenicity, highlighting its potential as an improved alternative to current clinical standards for hydrophobic implantable materials.

### Evaluating the Biocompatibility of Fatty Aid-Modified Silk

The cytocompatibility and hemocompatibility of FAsilk was evaluated to ensure that the chemical modification process did not adversely impact biocompatibility. Cytocompatibilty was evaluated through the ISO-993:5 protocol.^31^ Silk plastics and controls were incubated in cell culture media to extract potential toxins and leachants from the chemical modification. Mouse fibroblasts (L929) cultured to confluency were then exposed to the media extracts to assess cytotoxicity. As seen in Figure 5(a), cells exposed to media extracts from the native and FAsilk plastics showed comparable percent cell viability to media-treated controls, as well as the cells exposed to media extracts from polyurethane films (negative control). In contrast, cells exposed to media extract from zinc diethyldithiocarbamate (ZDEC) polyethelene films (positive control) showed significantly reduced viability. Ordinary one-way ANOVA showed statistically significant effect between treatment groups [F(5, 30) = 132.3, p < 0.0001]. Post-hoc pairwise Tukey tests showed no statistically significant difference between native silk, FAsilk and the negative control (Figure 5a). The statistically significant decrease in percent cell viability as compared to untreated samples was attributed to the effect of cells exposed to fresh media in the untreated control throughout the experiment. This was further supported by Live/Dead imaging (Figure 5(b)) where cells exposed to native and FAsilk extracts showed similar numbers of green (live) cells and comparable cell morphologies to the negative controls.

**Figure 5.**
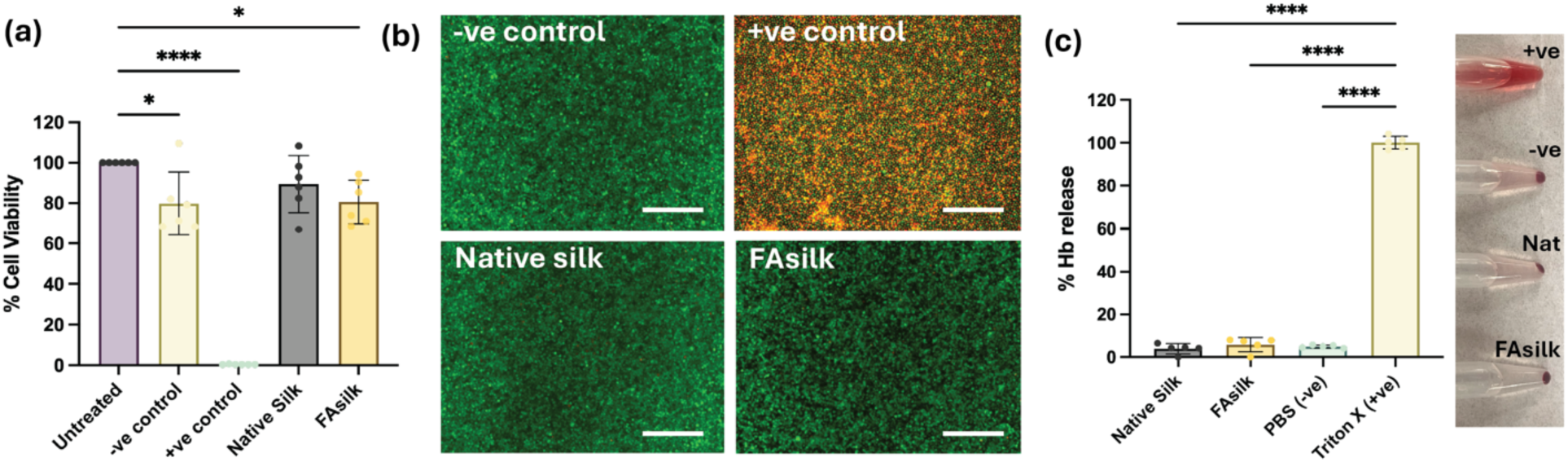
*In vitro* biocompatibility studies. (a) %Cell viability tested using media extracts of Native and FAsilk as well as negative and positive control substrates and untreated cells. %Cell Viability was normalized against untreated control. (b) Live/Dead staining and imaging of media-treated L929 cells where similar (green) cell morphologies are observed on -ve films as well as Native and FAsilk. Clear dead cells (red) are visible on +ve control. Scale bar is 200 μm. (c) Hemolysis assay conducted by treating red blood cells extracted from whole human blood with media extracts. High %Hb release (as seen in the positive control) corresponds with lysed RBCs indicative of poor hemocompatibility. Both silk samples exhibit similarly low hemolysis to PBS and exhibit a pellet of whole RBCs upon centrifugation. Asterisks indicate adjusted p-values. p < 0.05 (*) and p < 0.0001 (****). Unmarked comparisons are not significant.

Hemocompatibility of the modified silk materials was assessed similarly using media extracts, following the ISO-993:4 protocol.^32^ Media extracts were prepared by incubating sterilized silk samples and controls in PBS and mixed in equal volumes with a 1% RBC solution. Equal volumes of (500 μL each) both the media extracts and 1% RBC solutions were mixed and incubated at 37°C for 1 h. Solutions were then centrifuged to allow intact RBCs to settle and the absorbance of the supernatant was measured at 405 nm. The percent lysis was calculated by normalizing against samples containing deionized water (positive control), which was assumed at 100% lysis. As seen in Figure 5(c) FAsilk samples showed minimal lysis of the RBCs, comparable with native silk and the negative control, indicating that the chemical modification was hemocompatible. Ordinary one-way ANOVA showed statistically significant effect between treatment groups [F(4,20)= 1679, p <0.0001]. However, post-hoc pairwise Tukey tests only showed statistically significant differences with the triton-X positive control, as indicated in Figure 5(c).

Biocompatibility was evaluated *in vivo* following subcutaneous implantation in mice. Native and FAsilk plastics were subcutaneously implanted in the back of the mice for 7 days. Additionally, a group of untreated mice were included. All animals were monitored daily for body weight, signs of distress, and food and water intake. Figure 6(a) illustrates the timeline of the animal study. As seen in Figure 6(b), no major changes in weight were recorded in mice across the three groups. Two-way RM (repeated measures) ANOVA showed statistically significant effects of days post surgery [F(6,72) = 54.30, p < 0.0001] and the interaction of days post-surgery and treatment on the weight of the mice [F(12,72) = 4.965, p < 0.0001]. Post-hoc pairwise Tukey HSD test showed statistically significant lowering of weight in the surgical groups (native and FAsilk) on days 2 and 3 post-surgery, as compared to untreated controls. This effect was attributed to the treatment groups recovering from surgery in the beginning.^33^ As seen in Figure 6(b) the body weight of mice in both treatment groups recovered over time, exhibiting no statistically significant differences by day 7.

**Figure 6.**
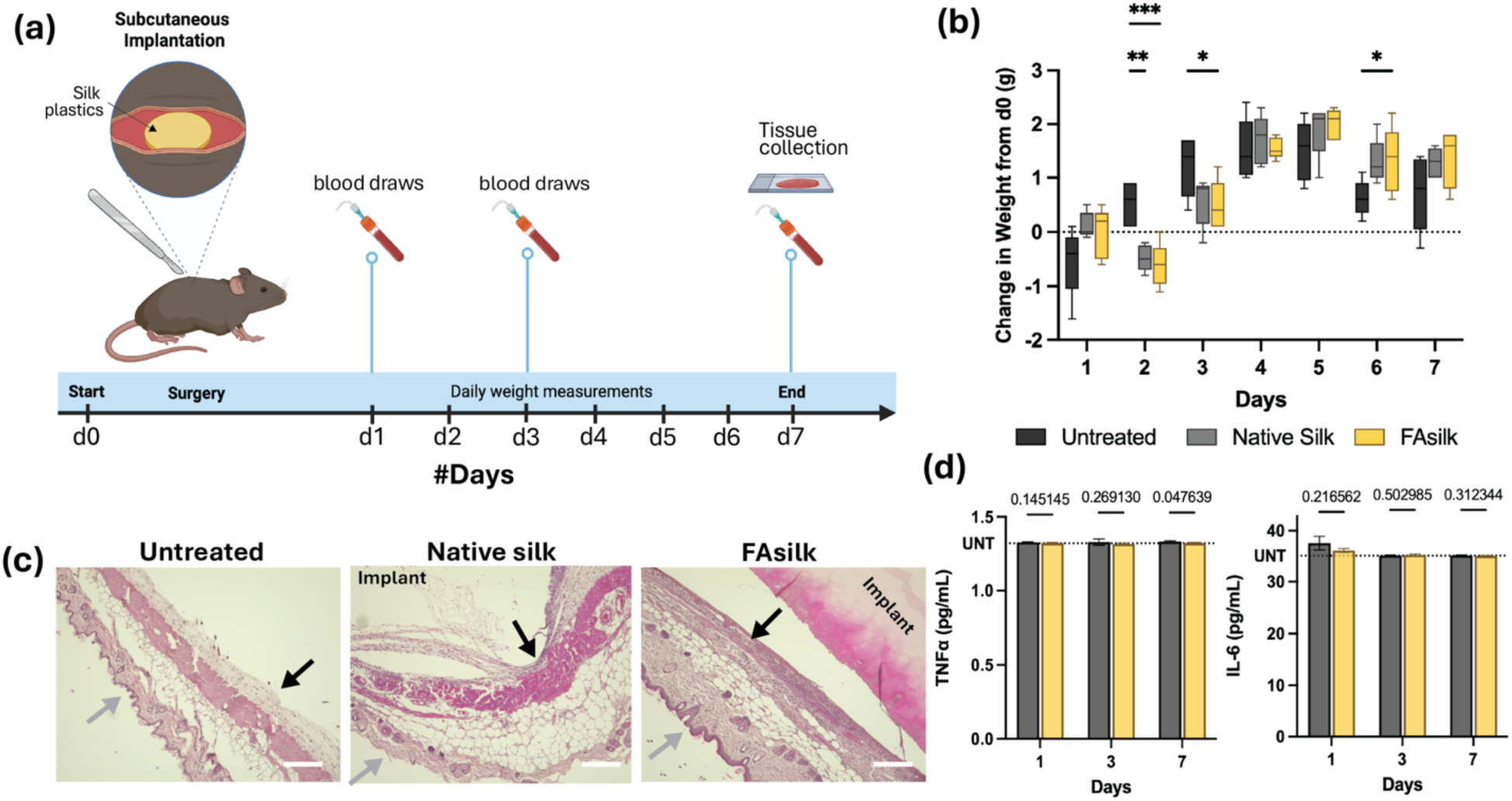
*In vivo* biocompatibility study. (a) Schematic depicting timeline of animal study. Mice were subcutaneously implanted with native or FAsilk plastics on day 0 and sacrificed on day 7. Weight was recorded daily and blood draws were conducted on days 1,3 and 7. Tissue from the implantation region were extracted on day 7. (b) Change in body weight of mice normalized to their weights prior to surgery on day 0. No significant weight loss was observed between mice of treatment (native, FAsilk) and untreated groups. (c) H&E stained images of tissue samples collected from implantation area. Gray arrows indicate the skin layer while black arrows indicate the layer interacting with implant. High cell infiltration (purple) observed in native and FAsilk implants. (d) Scale bar is 200 μm. Statistical analysis was conducted via two-way ANOVA analysis with n = 5 per treatment group. Asterisks indicate adjusted p-values. p < 0.05 (*), p < 0.01 (**), and p < 0.001 (***). Unmarked comparisons are not significant.

Histological analysis was conducted on skin tissue samples extracted from the implantation site on day 7, followed by hematoxylin and eosin (H&E) staining. Fixed and stained tissue samples were imaged via brightfield optical microscopy and the images are shown in Figure 6(c) as well as Figure S8, where additional images and lower magnification images are included. Peri-implant cellular accumulation was observed at the implant site in the case of both the native and FAsilk plastics, as compared to the untreated groups. This effect is consistent with acute localized inflammatory response surrounding dense implant material post-surgery and does not indicate poor biocompatibility, especially as native silk implants exhibit similar behavior and have demonstrated long-term biocompatibility. This was further confirmed by cytokine levels (TNFα, IL-6) detected in animal serum serum on days 1, 3 and 7 post surgery via multiplex immunoassays (Figure 6d). Both, native and FAsilks showed a slightly increased in IL-6 level on day 1 which stabilized to normal levels on subsequent days. There was no significant change in TNFα levels across the 7 days post-surgery across all 3 groups. Furthermore, no significant differences were observed between the native and FAsilk groups on any day post-surgery, determined by unpaired t-tests with Welch’s correction and adjustment for multiple comparisons using the two-stage step-up Benjamini, Krieger, and Yekutieli method to control the false discovery rate. Adjusted p-values are reported in Figure 6(d). Therefore, fatty acid modification did not adversely impact biocompatibility and FAsilk materials showed comparable biocompatiubility to the native silk materials.

## Conclusions

This study establishes a fatty acid functionalization approach for transforming silk into a water-stable, bioresorbable, hydrophobic, implantable biomaterial. Using an anhydrous esterification strategy enabled with ionic liquids, fatty acids were covalently linked to the silk backbone. FAsilk retained thermalmolding into dense plastic materials, despite significant changes to the protein structure, likely due to the hydrophobic interactions between the fatty acid groups. This enabled FAsilk to be thermoplastically molded and fabricated for implants. These FAsilk plastics exhibited substantial resistance to aqueous and enzymatic degradation when compared to the control, unmodified silk samples, preserving structural and mechanical integrity and overcoming a key limitation of conventional silk-based materials due to swelling in water and loss of some mechanical integrity. The material characteristics were accompanied by compatible interactions with biological milieus. FAsilk reduced microbial contamination and macrophage adhesion relative to native silk, and minimized blood clot formation, indicating reduced potential for thrombosis, foreign body response and infections. Furthermore, FAsilk maintained cytocompatibility and hemocompatibility, and subcutaneous implantation showed minimal proinflammatory responses or adverse impact to animal health. Together, these findings demonstrate that fatty acid functionalization can impart hydrophobicity and aqueous stability to silk materials while preserving biodegradability, processability and biological compatibility. FAsilk provides a bioresorbable alternative to synthetic hydrophobic materials for implants, including medical devices, tissue-engineered grafts and drug delivery systems, by balancing material stability with biological integration and function.

## Methods

### Synthesis and Fabrication of Fatty Acid-modified Silk Materials

#### Materials

All chemicals, reagents and solvents were purchased from MilliporeSigma (Burlington, MA, USA) unless otherwise specified. *Bombyx mori* silk was purchased from Sofregen Medical (Framingham, MA, USA)

#### Silk preparation

Silk fibers were degummed as previously described.^34^ Briefly, *Bombyx mori* silk cocoons were cut and boiled in 0.02M Na₂CO₃ solution for 30 min, then rinsed three times in dH₂O (deionized water) for 10 min per rinse and subsequently dried overnight at RT and stored on the benchtop until further processing. To prepare aqueous silk solutions, dried silk was dissolved in 9.3M LiBr at 60*°*C for 3 h, then dialyzed against diH₂O for 72 h with 6 water changes. Following dialysis, the solution was filtered through a 40 µm cell strainer and centrifuged (9,000 rpm for 20 min) to remove any insoluble particulates. The solution was then diluted to a 1% w/v working concentration and frozen overnight at -80*°*C before lyophilization at -82*°*C, 0.006 bar for 3 days. Lyophilized powders were stored at -20*°*C until plasticization.

#### Preparing silk fibroin ionic liquid for fatty acid modification

To generate an ionic liquid (SF_IL_), 1.0 g of degummed silk was melted in 7.0 g of 1-butyl-3-methylimidazolium chloride (BMIM-Cl) at 100*°*C for 3 h. The molten liquid was then diluted with 15 mL of N,N-dimethylformamide (DMF), with 1.8 g of urea added when applicable. The reaction mixture was stirred until homogenous, followed by cooling to RT.

#### Chemical modification of fatty acids on SF_IL_

To the SF_IL_ prepared above, 2.0 mL of linolenic acid, 1.0 g of 1-ethyl-3-(3-dimethylaminopropyl)carbodiimide (EDC), and 1.0 g of 4-dimethylaminopyridine (DMAP) were sequentially added, with complete dissolution of each reagent before the next addition. To minimize moisture ingress and light-induced fatty acid oxidation, the reaction vessel was sealed with Parafilm and wrapped in aluminum foil before overnight stirring at RT in the fume hood. The following day, the mixture was transferred to a 3.5-kDa MWCO dialysis membrane and dialyzed against dH₂O for 72 h with 6 water changes. After dialysis, the precipitates (FASilk) were washed for an additional 72 h in dH₂O with 2 water changes daily, followed by final centrifugation (9000 rpm for 20 min) before collection for lyophilization. FAsilk precipitates were frozen overnight at -80*°*C before lyophilization at -82*°*C, 0.006 bar for 3 days. Lyophilized powders were stored at -20*°*C until plasticization.

#### Fabrication of silk thermoplastics

Lyophilized FAsilk and native silk powders (80 mg per plastic) were loaded into pre-designed circular stainless steel molds (5.5, 10, and 12 mm diameter) and thermomechanically compressed at variable temperatures and 540 psi for 15 min following our established methods.^10, 11^

### Characterization of FAsilk

#### NMR Characterization of chemically modified silk

To structurally characterize the modified silk formulations, lyophilized powders of native silk, SF_IL_ (silk fibroin ionic liquid), FASilks and free fatty acids were dissolved in deuterated dimethyl sulfoxide (DMSO-D_6_) at a concentration of 5-8 mg/mL (depending on solubility). ^1^H-NMR Spectroscopy was performed with a Bruker Avance Neo 400 (Billerica, MA, USA).

#### ^1^H and ^13^C solid-state NMR of native and FAsilks

ssNMR were performed in a 4mm broadband CP/MAS (cross-polarization/magic angle spinning) probe on a Bruker Avance III 600 MHz NMR spectrometer at ambient temperature. The following experiments were conducted: (1) ^1^H single-pulse NMR experiment, which was conducted without magic angle spinning (MAS). Recycle delay was 6 s. Dwell time was 2 μs. (2) Cross polarization (CP) experiments. Contact time was 2 ms and recycle delay was 4 s. (3) Direct polarization (DP) experiments with high power 1H decoupling. Recycle delay was 2 s so that only signals from the most mobile segments were collected. (4) ^1^H spin diffusion experiment. Dephasing time was 26 μs and contact time was 2 ms. Magic angle spinning (MAS) speed was 8 kHz for all MAS experiments.

#### ATR-FTIR characterization of silk powders and plastics

ATR-FTIR spectroscopy was conducted using a JASCO FTIR 6200 spectrometer (JASCO, Tokyo, Japan) with a MIRacle attenuated total reflectance (ATR) with germanium crystal on powders of native silk, SF_IL_ and FAsilks as well as plastics of native and FAsilks. The measurements were performed by averaging 64 scans with a resolution of 4 cm^−1^ within the 4000-600 cm^-1^ wavenumber range. For each sample, FTIR spectra of n=3 replicates were measured. Spectra were baseline corrected and deconvoluted, and % β-sheet content (1616-1637 cm-1), random coil (1638-1655 cm-1), α-helix (1656-1662 cm-1), and β-turns (1663-1696 cm-1)) were calculated using the Fourier self-deconvolution method (Lorentz deconvolution) using Jupiter notebooks. Data was plotted and statistical analysis was conducted using GraphPad Prism 10 via Ordinary two-way ANOVA with Tukey’s multiple comparisons tests and f-values, adjusted p-values and degrees of freedom (DF) were reported.

#### Flexural modulus via three-point bending

Mechanical properties of native and FASilks were assessed via three-point bend test on an Illinois Tool Works Instron (Norwood, MA, USA) Model 3366, equipped with a load cell of 100 N. Plastics were prepared by thermomechanical compression at 125*°*C, 540 psi for 15 min into circular discs, and evaluated in both dry and hydrated (post 7-day incubation in PBS) states. Samples (n= 3 per condition) were placed onto two support points with a span of 6mm and a load was applied at a rate of 0.1 mm/min until break or 2mm total deflection, for dry and hydrated samples respectively. Flexural modulus (kPa) was calculated by measuring the slope of the linear regions of resultant force-displacement curves. % Strength retained was calculated for each wet sample by normalizing against the corresponding average dry flexural strength. Statistical analysis was conducted using Graphpad Prism 10 via Ordinary one-way ANOVA with Tukey’s multiple comparisons tests and tests and f-values, adjusted p-values and degrees of freedom (DF) were reported.

#### Measuring surface roughness and 3D surface morphology

3D surface morphology imaging was conducted on dry and wet samples using Keyence Surface Profiler VK-X4000 (Keyence America, Itasca, IL, USA). Surface roughness was estimated using the surface roughness program in the manufacturer’s software. Average surface roughness was calculated with n = 3 measurements per sample. Statistical analysis was conducted using Graphpad Prism 10 via Ordinary one-way ANOVA with Tukey’s multiple comparisons tests and tests and f-values, adjusted p-values and degrees of freedom (DF) were reported.

#### Thermogravimetric analysis

Thermal decomposition properties of silk powders were evaluated using a Texas Instruments Q500 thermogravimetric analyzer (TA Instruments, New castle, DE, USA). For each sample, 5 to 7 mg of powder were weighted and encased in aluminum sample pans with vent holes punched. Samples were heated at 10°C/min from room temperature to 600°C. Nitrogen gas was supplied to the chamber with a balance purge flow rate of 40 mL/min and a sample purge flow rate of 50 mL/min.

### Stability Testing for Silk

#### Aqueous stability

FASilk and native SF plastics were plated in quadruplicate (n=4 technical replicates per condition) in 24-well plates. Initial dry weights were recorded before immersion in 1.0 mL of DPBS, followed by incubation at RT for 7 days. During immersion, the wet weight of samples were recorded at regular intervals (time points: 1 h, 24 h, 48 h, 96 h, and 168 h). Prior to each measurement, excess water moisture was removed using compressed air. Aqueous stability was quantified by calculating %water uptake relative to the initial weight for each sample. Statistical analysis was conducted using Graphpad Prism 10 via Ordinary one-way ANOVA with Tukey’s multiple comparisons tests and tests and f-values, adjusted p-values and degrees of freedom (DF) were reported.

#### Enzymatic stability

Native and FASilk plastics were prepared in triplicate (n=3 technical replicates per condition) in a 24-well plate. Initial masses were recorded prior to immersion in 1.0 mL of Protease XIV Solution (prepared in PBS) and incubation at 37*°*C. Samples were incubated in 10 U/mL Protease XIV solution the first 12 days and in 20 U/mL Protease XIV solution for the remainder of the assay. Prior to each measurement, excess moisture was removed using a stream of compressed air before weighing. Samples were then re-submerged in protease solution and returned to the incubator until the following time point. Incubated protease solutions were fully replaced every 48 h with freshly prepared solutions. Enzymatic stability of each sample was characterized by calculating mass change and reduced weight of each sample at every timepoint relative to the initial measurement. Statistical analysis was conducted using Graphpad Prism 10 via Ordinary one-way ANOVA with Tukey’s multiple comparisons tests and tests and f-values, adjusted p-values and degrees of freedom (DF) were reported.

### Evaluating the Thrombogenicity of FAsilk

Human whole peripheral blood was purchased from StemCell technologies (Cambridge, MA, USA) and collected from a 30 year old healthy female in EDTA and shipped one week prior to experiments.

#### Material preparation

Biomaterial thrombogenicity was assessed using a whole blood clotting assay adapted from established protocols. Experimental conditions include native silk, FASilks, PTFE (clinical benchmark for blood-contacting materials), TCP, as well as dH₂O which served as a control for hemolysis and hemoglobin (Hb) release. All plastics were sterilized with ethylene oxide (ETO) before individual placement in a 24-well plate (n=3 plastics per condition). dH₂O wells were filled with 500 µL of sterile ultrapure water.

#### Blood clotting assay

Human whole blood vials were inspected for premature coagulation prior to use and samples were tested using established protocols.^27^ 10 µL of blood was pipetted onto each surface and incubated at RT for 30 min. dH₂O wells were not subjected to the incubation period but the blood was simply mixed into the well. Following incubation, samples were transferred to new wells in a 24-well plate containing 500 µL of dH₂O to completely submerge the samples. Care was taken to not disturb the blood on samples. Samples were shaken at 100 rpm for 30 sec, then allowed to rest at RT for 5 min to release free Hb from unattached RBCs. For each well, 100 uL of the surrounding media was transferred to a 96-well plate (n=3 replicates per plastic) and absorbance measured at 540 nm. All values were normalized to the dH₂O control, which was defined as 100% Hb release. Thrombogenicity was characterized by assessing Hb release (%) across the experimental conditions and lower % Hb release corresponded to high clot formation. Average across each sample was first obtained and then averaged across a single experimental condition. Statistical analysis was conducted using Graphpad Prism 10 via Ordinary one-way ANOVA with Tukey’s multiple comparisons tests and tests and f-values, adjusted p-values and degrees of freedom (DF) were reported.

### Evaluating Microbial Adhesion

#### Bacterial cell culture

Freeze-dried stocks of *Staphylococcus aureus* were purchased from ATCC (Manassas, VA, USA) and tryptic soy broth and agar plates were obtained from BD Biosciences (Milpitas, CA, USA) and prepared and sterilized according to manufacturer’s protocol. To prepare streak plates, freeze-dried *S.aureus* was suspended in 5.0 mL of Tryptic Soy Broth (TSB) and incubated overnight at 37*°*C, 350 rpm in a shaking incubator. Bacteria was then streaked onto TSA plates and incubated without agitation at 37*°*C for 24 h. The following day, plates were inspected for isolated colonies, then sealed with Parafilm and stored at 4*°*C until use. To prepare cultures for seeding, isolated colonies from stored TSA plates were inoculated in 3.0 mL TSB and shaken overnight at 37*°*C, 350 rpm. The following day, cultures were washed twice in sterile 0.85% NaCl and centrifuged twice at 4000 rpm for 5 min each. The pellet was re-suspended in 2.0 mL PBS and concentration measured by optical density (1.0 OD_600_ = 10^9^ cfu/mL). A working bacterial seeding suspension was prepared by diluting the culture to 0.1 OD in TSB.

#### Biofilm formation study

Native SF and FASilk plastics were placed in a 6-well plate (n=1 plastic per condition) and washed in 2.0 mL sterile ultrapure dH₂O at RT for 3 h. After initial wash, dH₂O was replaced with 3.0 mL TSB, and the plastics were incubated at 37*°*C for 24 h for conditioning. The following day, plastics were transferred to a new 12-well plate, and 1.5 mL of the working bacterial suspension was seeded onto each surface before incubation at 37*°*C for 24 h without agitation to induce biofilm formation. TCP was included as the negative growth control.

#### Live/Dead Imaging

After 24 h, the plate was washed with 0.85% NaCl and stained with Baclight LIVE/DEAD bacterial cell viability kit, per manufacturer’s protocol. Staining solutions were prepared in 0.85% NaCl and kept in the dark until addition, after which samples incubated for 10 min at RT prior to imaging using a Keyence BZ-X700 fluorescence microscope with visible green and red filter cubes under 10X and 20X magnification.

### Evaluating Monocyte Activation and Macrophage Adhesion

#### Monocyte cell culture

THP1-ASC-GFP cells were acquired from InvivoGen (San Diego, CA, USA) and cultured in RPMI media (Gibco, Thermo Fisher, Waltham, MA, USA), supplemented with 10% heat-inactivated Fetal Bovine Serum (FBS, from Gibco) and 1% Antibiotic-Antimycotic (Gibco). Cells were cultured to confluency at 37*°*C, 5% CO₂, and passaged at least once before experimental use.

#### Monocyte activation test

THP1ASC-GFP cells were centrifuged at 300 x g for 5 min to obtain pellet and resuspended at 10^5^ cells/ mL density. ETO-sterilized native and FAsilk samples were first conditioned in culture media at 37 °C for 24 h and then placed in each well of a 24-well plate. Cells were seeded on top of each sample by placing 1 mL of seeding solution per well and incubated at 37*°*C, 5% CO₂ for 3 h. Cells were seeded with media containing LPS (lipopolysaccharide) as a positive control. After 3h 10 μL of DAPI solution (Thermofisher) was directly added to each well to stain cell nuclei.

#### Monocyte differentiation and macrophage adhesion

THP-1 cells were treated with phorbol 12-myristate 13-acetate (PMA) at a concentration of ∼14 ng/mL in 7.0 mL of culture media, then incubated for 24 h at 37*°*C, 5% CO₂ to induce macrophage-like phenotype. Following incubation, cells were manually scraped, centrifuged at 300 x g for 5 min, and re-suspended in PMA-free media for seeding.

Prior to cell seeding, native and FASilk plastics were ETO-sterilized and placed in a 24-well plate. Materials were conditioned in 1.0 mL of complete media for 24 h at 37*°*C, 5% CO₂. Following material conditioning, PMA-treated macrophage-like THP1 cells were seeded onto the material surfaces at 1 x 10^5^ cells/well and incubated for 24 h at 37*°*C. TCP was tested as negative control. After cell attachment, the wells were spiked with LPS at a final concentration of 1.0 µg/mL per well and incubated for a final 24 h at 37*°*C.

#### Imaging

Cells were imaged using a Keyence BZ-X700 fluorescence microscope with visible green, blue and red filter cubes.Three images were acquired per sample condition across Brightfield, DAPI, and GFP channels using 4X and 10X magnification.

### Evaluating cytocompatibility

#### Cell culture

L929 murine fibroblasts were purchased from ATCC and cultured in Dulbecco’s Modified Eagle Medium (DMEM, Gibco), supplemented with 10% Fetal Bovine Serum (FBS, Gibco) and 1% Antibiotic-Antimycotic (Gibco). Cells were regularly passaged once weekly after reaching ∼90% confluency, with media changes every 3-4 days. Cells were detached with 0.25% trypsin EDTA (Gibco) and centrifuged at 300 x g for 5 min.

#### Preparation of media extracts

ETO-sterilized FASilk plastics and purchased experimental controls—polyurethane (-ve control) and zinc diethyldithiocarbamate (ZDEC) polyethylene (+ve control) —were deposited in 6-well plates for leachate generation (n=3 plastics per condition per well), per ISO-993:5 protocol.^31^ Samples were submerged in 2.0 mL of sterile ultrapure dH₂O and incubated at 37*°*C, 5% CO₂ for 24 h for an initial wash, then replaced with 3.0 mL of complete medium and further incubated at 37*°*C for an additional 24 h for leachate collection.

#### % Cell viability assessment via PrestoBlue assay

Cytotoxicity assays were performed in accordance with the ISO 993-5 protocol (International Standardization Organization 10 993: Biological Evaluation of Medical Devices, Part 5: Tests for Cytotoxicity: *in vitro* methods).^31^ L929 murine fibroblasts were seeded in 96-well plates at a density of 1.0 x 10^5^ cells/well and incubated at 37*°*C, 5% CO₂ for 48 h prior to treatment. Culture medium was then replaced with leachate solutions (n=8 replicates per condition) and incubated for an additional 48 h before cell viability quantification with PrestoBlue, per manufacturer’s instructions (Invitrogen). A growth control condition was also included, containing only the seeded cell suspension in complete growth medium. % Cell viability for each condition was calculated against untreated growth control. Statistical analysis was conducted using Graphpad Prism 10 via Ordinary one-way ANOVA with Tukey’s multiple comparisons tests and tests and f-values, adjusted p-values and degrees of freedom (DF) were reported.

#### Live/ Dead Imaging

Two wells from the above experiment were stained with Live/Dead Cell staining kit (Invitrogen) according to manufacturer’s protocol and images using Keyence BZ-X700 Fluorescence microscope where red channel represents dead cells and green channel represents live cells.

### Evaluating hemocompatibility

#### Preparation of RBCs

Human whole peripheral blood was purchased from StemCell technologies, collected from a 30 year old healthy female in EDTA, shipped one week prior to experiments, and assessed for premature coagulation prior to experimental use. Vial contents were first transferred to a falcon tube, centrifuged at 1700 x g for 5 min, then re-suspended in PBS twice for initial wash. Red blood cells were counted, then diluted 1:100 to obtain a working 1% suspension (∼2.5 x 10^5^ cells/mL).

#### Preparation of media extracts

Sterilized native and FASilk plastics were plated in a 12-well plate (n=2 plastics per well per condition), washed with 2.0 mL of ultrapure dH₂O for 2 h before incubation in PBS for 24 h at RT.

#### Hemolysis assay

Assay was performed in accordance with ISO 993-4 (International Standardization Organization 10 993: Biological Evaluation of Medical Devices, Part 4: Selection of Tests for Interactions with Blood), following established protocols.^32^ After media extract generation, 50 µL of each leachate solution was mixed with 50 µL of 1% RBC suspension and pipetted into a 96-well PCR plate (n=6 replicates per condition). Controls were set up in parallel, with PBS as negative control and 10% v/v Triton X-100 as positive control. The plate was sealed and incubated at 37*°*C, 5% CO₂ for 3 h, then centrifuged twice at 2800 rpm for 1 min. Following centrifugation, 70 µL of each supernatant was carefully transferred to a new flat, transparent bottom 96-well plate without disturbing the pellet and absorbance was measured at 405 nm using a microplate reader. % hemolysis was calculated by normalizing sample absorbances to the positive control assumed at 100% hemolysis. Statistical analysis was conducted using Graphpad Prism 10 via Ordinary one-way ANOVA with Tukey’s multiple comparisons tests and tests and f-values, adjusted p-values and degrees of freedom (DF) were reported.

### Biocompatibility study

All animal procedures were performed with ethical compliance and approval by the Institutional Animal Care and Use Committee at Tufts University. Lyophilized native SF and FASilk powders were thermomechanically compressed in circular 5.5 mm diameter molds at 125*°*C, 90 psi for 15 min to yield plastics of appropriate dimensions for subcutaneous implantation. Samples were washed in PBS for 24 h at RT before ETO-sterilization (n = 5 plastics per condition). Adult female C57BL6 mice approximately weighing 20 g each (n = 5 per treatment group) were purchased from Charles River Laboratories (Cambridge, MA, USA) and acclimated prior to surgery. All surgical equipment was sterilized prior to surgery using autoclave and re-sterilized during surgery using a hot glass bead sterilizer at 300 °C. On day 0 (surgery day), initial weights were recorded and mice were shaved. 30 min prior to surgery, mice were administered with a single subcutaneous injection of 1 mg/ kg of sustained-release buprenorphine for pain management. Mice were anesthetized with isofluorane throughout surgery. Surgical site (back) was cleaned using alcohol prep pads (3 times) and disinfected using Dynarex 10% povidone-iodine swabs (3 times). A ∼10 mm subcutaneous incision was made on the back and native or FAsilk plastics (5 mm each) were implanted in each mouse. Surgical incision was closed using surgical clips. One group of untreated mouse (n = 5) were included along with treatment groups. Mice health and body weight was monitored daily for 7 days. On day 7, mice were euthanized and tissue surrounding implants was collected for histopathological analysis. Statistical analysis of changes in body weight was conducted using Graphpad Prism 10 via repeated measures two-way ANOVA with Tukey’s multiple comparisons tests and and f-values, adjusted p-values and degrees of freedom (DF) were reported.

#### Histology

Mouse skin samples were collected from the implantation sites and fixed in 4% paraformaldehyde (PFA, Thermo Fisher) at 4°C until processing. Samples were then submitted to the Tufts Comprehensive Histology and Pathology Core Services for embedding, sectioning, and hematoxylin and eosin (H&E) staining according to standard protocols. Samples were imaged using a Keyence BZ X710 fluorescence microscope using brightfield.

#### Multiplex immunoassay

Serum was collected on days 1, 3, and 7. Samples were preserved at -20°C until the assay was performed. A Luminex Discovery Assay (R&D Systems, Minneapolis, MN, USA) that included IL-6 and TNFα was conducted on serum samples diluted 1:2 and following the manufacturer’s protocol. Samples were run on a Luminex MAGPIX Multiplexing system, Data were collected using Luminex xPONENT software, exported to a .CSV and analysed using Excel and GraphPad Prism.

### Statistical information

All statistical analysis were conducted using GraphPad Prism 10 via ANOVA (ordinary oney-way, ordinary two-way, or repeated-measures two-way) as reported within the relevant methods and results sections. Unless otherwise noted in the relevant methods or results section, at least n=3 replicates were tested for establishing statistical significance. Where appropriate, post hoc multiple-comparison tests were performed using Tukey’s honestly significant difference (HSD) test. Adjusted p values accounting for multiple comparisons were reported in the relevant results section.

## Supporting information

SI_revised

SI_original

## Data Availability

The authors declare that all the data supporting the findings of this paper are available within the paper and in the supplementary information document.

## Code Availability

ATR-FTIR data was processed and analyzed using Jupyter notebooks. Code for this is provided as a supplementary file.

## Acknowledgements

This work was supported by the National Institut of Biomedical Imaging and Bioengineering [NIH-P41EB027062] and the Department of War [FA9550-23-1-0606]. M.L.A. acknowledges support from the National Institute of General Medical Sciences for the Tufts IRACDA Program [NIH-K12GM133314].

## Author Contribution Statement

S.G. contributed to the conceptualization, methodology, investigation, validation, and analysis of all experiments, data visualization, and writing. E.Z., M.L.A., E.D.G., J.K.S., and S.M.S., contributed to investigation, validation, and analysis of several experiments. C. L. contributed to the methodology for animal studies and management of IACUC compliance. D.L.K. contributed to conceptualization, supervision, project administration and funding acquisition.

