## Supplementary material for "Molecular Engineering of Silk via Fatty Acid Esterification for Water-Stable, Anti-fouling, Biocompatible, Implantable Materials": SI_original

### Author Information

Sanjana Gopalakrishnan, Emilee Zhang, Mariah L. Arral, Edward B. Gordon, Sophie M. Steedly, Jugal K. Sahoo, Chunmei Li, David L. Kaplan*

Affiliation

**Department of Biomedical Engineering, School of Engineering, Tufts University, Medford, MA 02155**

Corresponding author

Correspondence to: Prof. David L. Kaplan

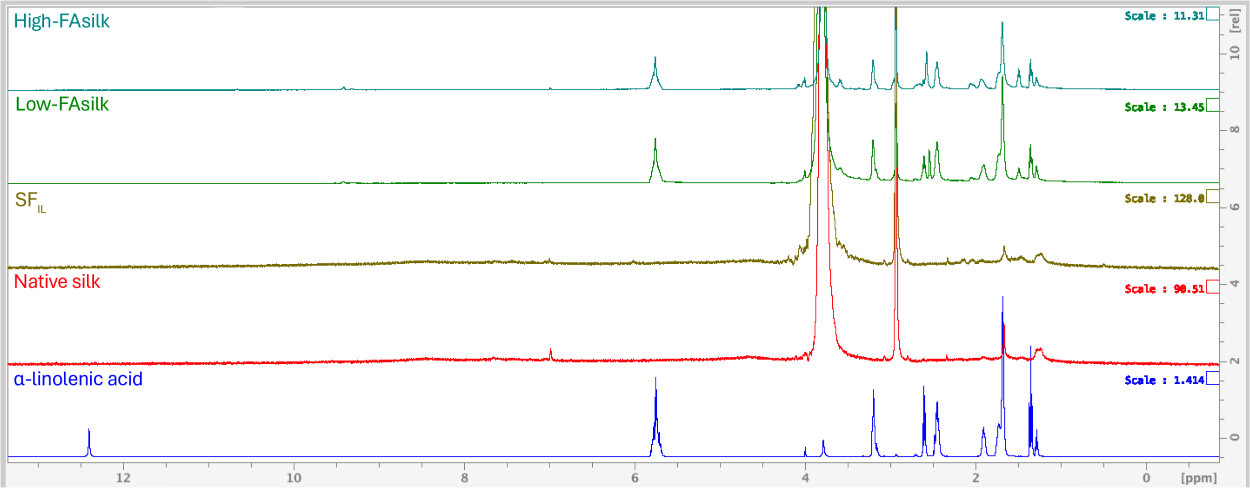


**Figure S1.** $^1$H NMR spectra showing Native silk, free α-linolenic acid and α-linolenic acid modified silk. Several α-linolenic acid peaks observed in the low-FAsilk and high-FAsilks consistent with modification. Black arrow indicates deshielded -COOH proton in free fatty acid which is absent in modified silks indicative of esterification.

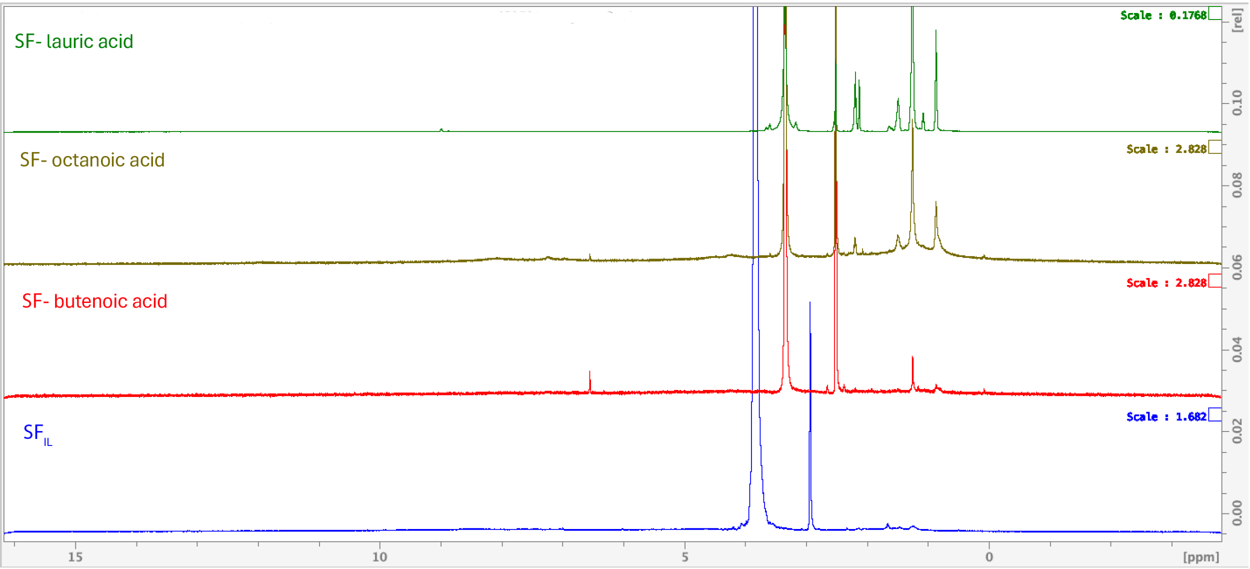


**Figure S2.** $^{1}$H NMR spectra showing other fatty acid modifications where, (a) is lauric acid-modified silk and free lauric acid, (b) is butenoic acid-modified FAsilk and free butanoic acid, and (c) is octanoic acid-modified silk and free octanoic acid.

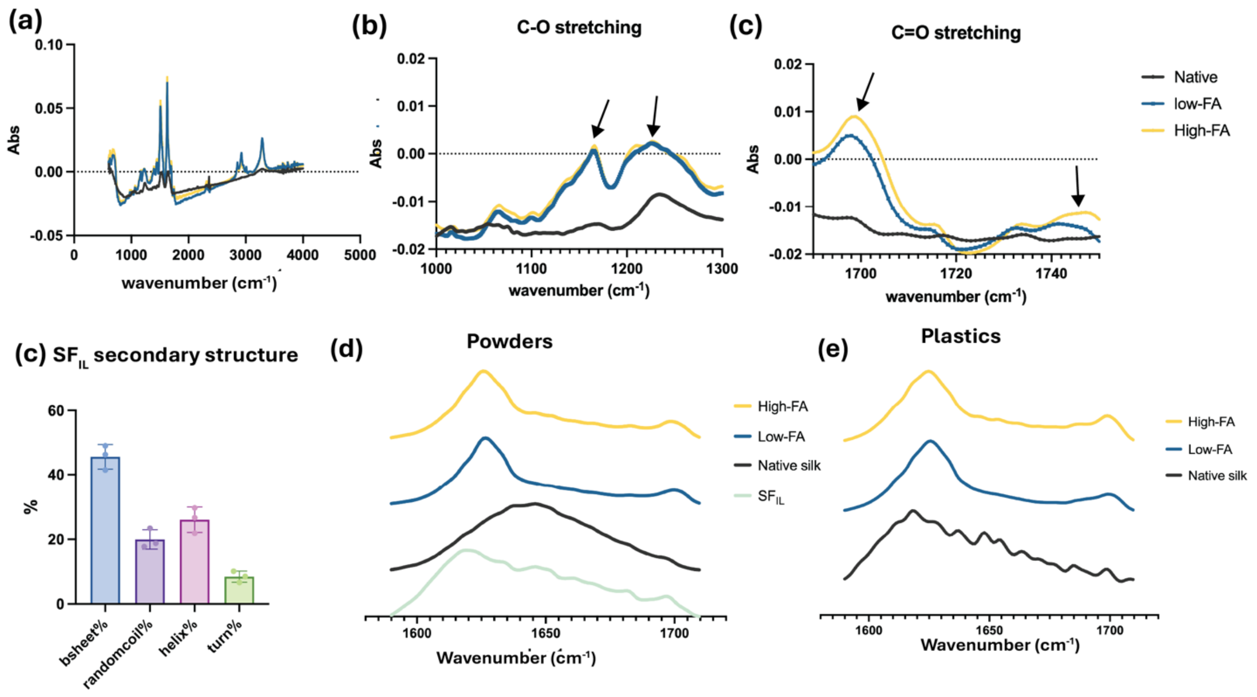


**Figure S3.** ATR-FTIR Spectra of Native silk, $SF_{IL}$, Low-FAsilk and High-FAsilk where (a) is the full ATR-FTIR spectra of lyophilized powders, (b) the spectra ranging from 1000-1300 $cm^{-1}$ showing the C-O stretching peaks in both low- and high-FAsilks, indicative of esterification, (c) the spectra ranging from 1690- 1750 $cm^{-1}$showing carboxyl stretching peaks at 1700 $cm^{-1}$ and 1745 $cm^{-1,}$ where a clear increase in peak intensity is observed in high-FA as compared to low-FA, consistent with higher degree of substitution. ATR-FTIR spectra used for deriving protein secondary structure of (a) silk powders and (b) silk plastics. Protein secondary structure is reported in Figure 2.

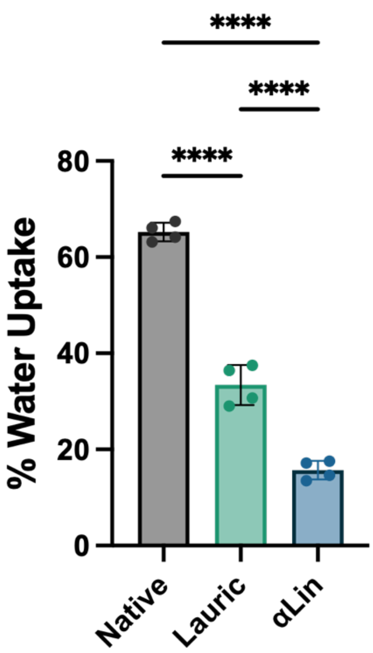


**Figure S4.** Water uptake studies conducted using fatty acid functionalized silk plastics modified with lauric acid (teal) and α-linolenic acid (blue), compared with unmodified native silk plastic (gray). Samples were incubated in PBS for up to 7 days. Both approaches show significant improvement to unmodified native silk however, α-linolenic acid modification outperformed lauric acid modification. Ordinary one-way ANOVA [$F(2,9) = 302.9$, $p<0.0001$] established statistically significant differences in %water uptake across type of modification. Post-hoc pairwise Tukey tests showed statistical significance across multiple pairs. Asterisks indicate adjusted p-values, where **** is $p<0.0001$.

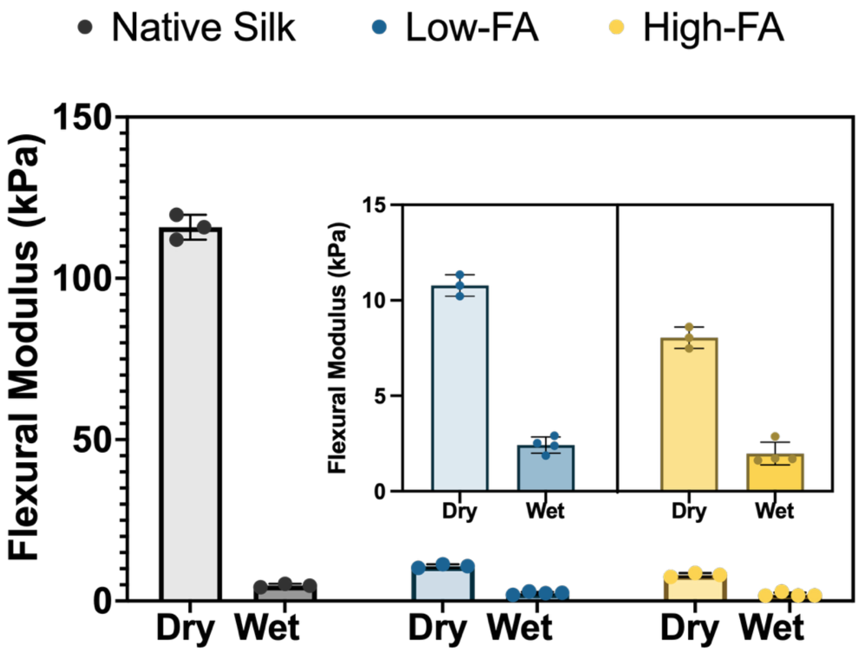


**Figure S5.** Flexural modulus (kPa) of dry and wet silk plastics. Inset shows Low-FA and High-FAsilk plastics.

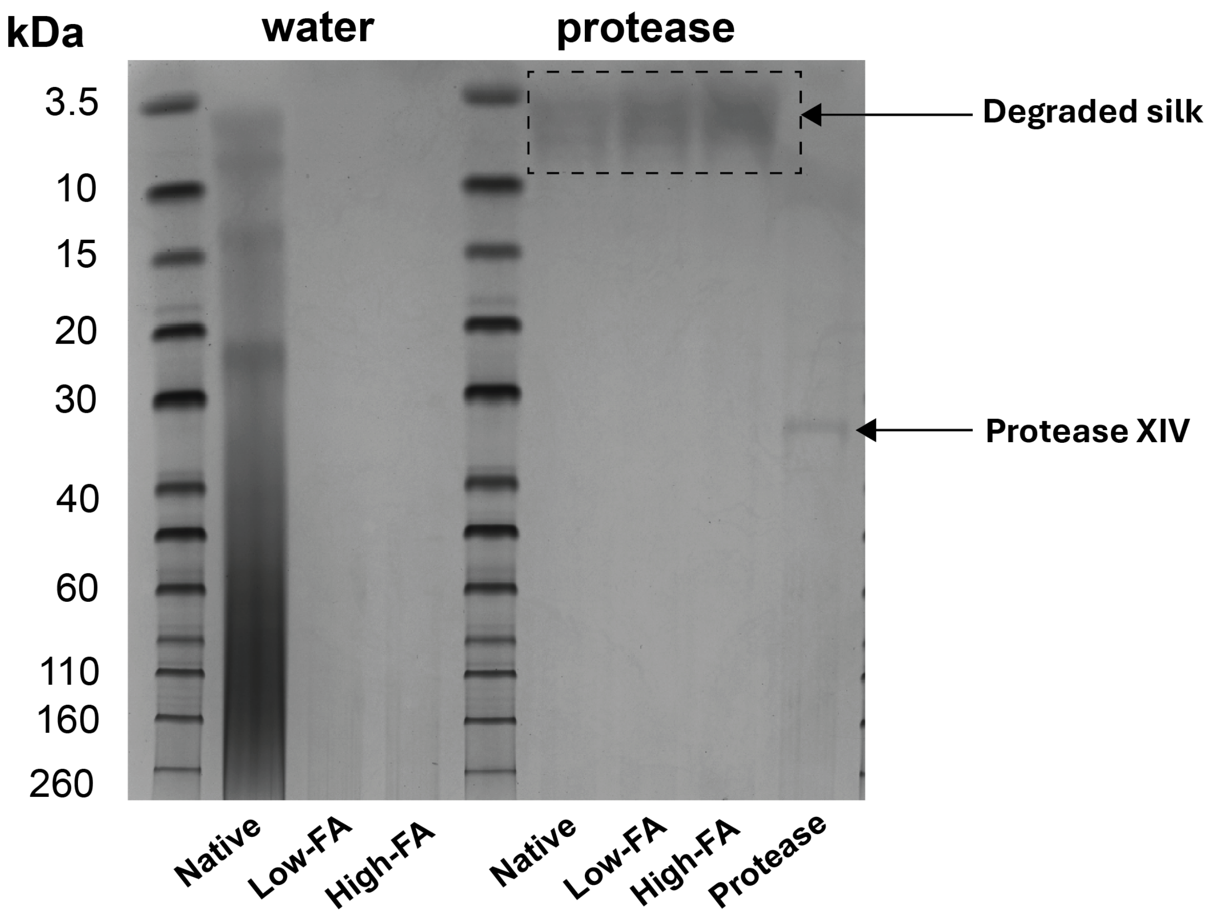


**Figure S6.** SDS-PAGE of native, low-FA and high-FAsilk solutions freshly prepared in deionized water (left) and post incubation in Protease XIV solution 1U/mL at 37 ºC for 24 h. Smear indicating a range of molecular weights observed for silk solution, while lighter bands observed for low-FA and high-FA due to poor solubility. In comparison, clear shift of all silk bands towards 3.5 kDa observed in protease indicating degradation.

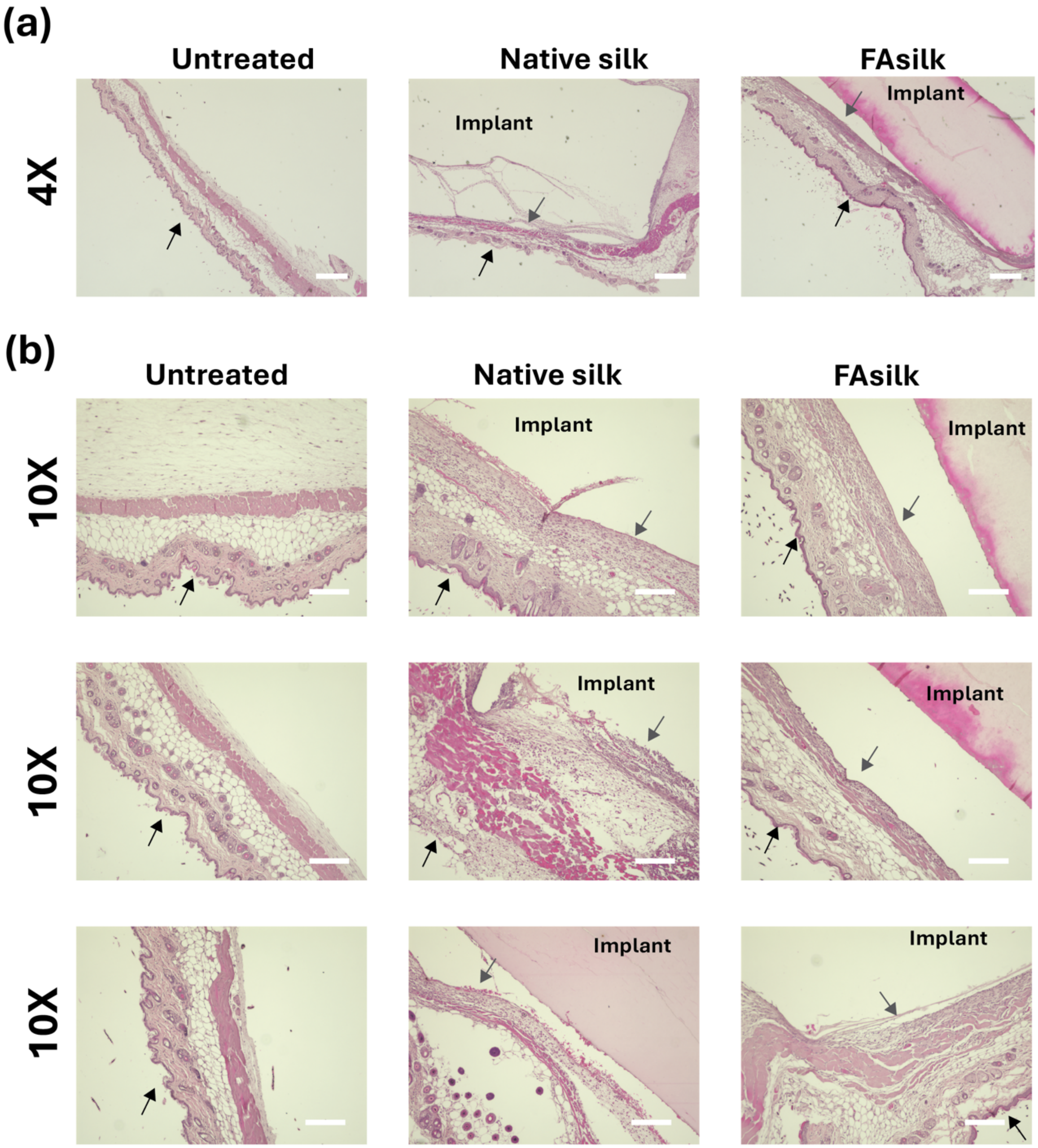


**Figure S7.** (a) 4X magnification of H&E-stained tissue samples and (b) Additional 10X magnification of H&E-stained tissue samples from untreated, native and FAsilk groups. Scale bar is 400 μm for (a) and 200 μm for (b). Black arrows represent skin and gray arrows show cell infiltration.
